# Cryo-EM structure of hexameric yeast Lon (PIM1) highlights importance of conserved structural elements

**DOI:** 10.1101/2021.11.24.469940

**Authors:** Jie Yang, Albert S. Song, R. Luke Wiseman, Gabriel C. Lander

**Author notes:** Authors contributed equally (Jie Yang, Albert S. Song).

## Abstract

Lon protease is a conserved ATP-dependent serine protease composed of an AAA+ domain that mechanically unfolds substrates and a serine protease domain that degrades unfolded substrates. In yeast, dysregulation of Lon protease (PIM1) attenuates lifespan and leads to gross mitochondrial morphologic perturbations. Although structures of bacterial and human Lon protease reveal a hexameric assembly, PIM1 was speculated to form a heptameric assembly, and is uniquely characterized by a ~50 residue insertion between the ATPase and protease domains. To understand the yeast-specific properties of PIM1, we determined a high-resolution cryo-EM structure of PIM1 in a substrate-translocating state. Here, we reveal that PIM1 forms a hexamer, conserved with that of bacterial and human Lon proteases, wherein the ATPase domains form a canonical closed spiral that enables pore loop residues to translocate substrate to the protease chamber. In the substrate-translocating state, PIM1 protease domains form a planar protease chamber in an active conformation and are uniquely characterized by a ~15 residue C-terminal extension. These additional C-terminal residues form an alpha-helix that is located along the base of the protease domain. Finally, we did not observe density for the yeast-specific insertion between the ATPase and protease domains, likely due to high conformational flexibility. Biochemical studies to investigate the insertion using constructs that truncated or replaced the insertion with a glycine-serine linker suggest that the yeast-specific insertion is dispensable for PIM1’s enzymatic function. Altogether, our structural and biochemical studies highlight unique components of PIM1 machinery and demonstrate evolutionary conservation of Lon protease function.

## Introduction

Protein homeostasis (proteostasis) – the dynamic balancing of protein composition in a cell – must be tightly controlled for cellular function and longevity^1^. As the hub of bioenergetic processes, mitochondria have an array of mechanisms to maintain proteostasis, one of which comprises proteases that site-selectively cleave substrates or degrade misfolded proteins^2^. These proteases acutely manage stress and prevent the accumulation of potentially toxic misfolded and aggregated proteins. However, as organisms age, the mechanisms that maintain proteostasis begin to fail – oxidative damage accumulates, healthy synthesis and degradation slows down – and hallmarks of age-associated pathology begin to emerge^3,4^.

In the mitochondrial matrix, the evolutionarily conserved Lon protease, or PIM1 in yeast, is essential for stress tolerance and proteostasis^5,6^. An adenosine triphosphatase associated with a variety of cellular activities (AAA+) serine protease, PIM1, modulates mitochondrial DNA copy number^7^ and maintains quality control and respiration^8^. Lon expression has been demonstrated to be modulated by various stressors, such as hypoxia and reactive oxygen species^9^. In yeast, the activity of PIM1 has been demonstrated to decline with age, and deletion of PIM1 shortens the replicative life span^10^, inhibits ATP-dependent degradation of matrix proteins, prevents growth on nonfermentable carbon source, and leads to gross mitochondrial morphologic perturbations^11^.

Despite the functional importance of PIM1 for mitochondrial proteostasis, an atomic model of the PIM1 oligomer is not yet defined. While initial negative stain electron microscopy of purified PIM1 combined with analytic centrifugation suggested that PIM1 is a heptamer^12^, atomic models of bacterial^13^ and human Lon^14^ present hexameric assemblies consistent with many AAA+ proteases^15^. Of note, PIM1 from *Saccharomyces cerevisiae* is defined by a unique insertion of approximately 50 highly charged residues between the ATPase and protease domains **(Supplementary Figure 1)**. Using cryo-electron microscopy (cryoEM), we aimed to define the oligomeric state of PIM1 from *S. cerevisiae* and understand the structural role of this charged insertion in protease enzymatic activity.

Here, we recombinantly expressed and purified *S. cerevisiae* PIM1 to reveal that PIM1 in the presence of translocating substrate indeed forms a right-handed spiral configuration consistent with orthologous Lon protease structures and other AAA+ proteases. Aromatic pore loop residues directly bind substrate at the central channel of the ATPase domain, indicating this protease functions through the canonical hand-over-hand substrate translocation mechanism that is coordinated with ATP binding and hydrolysis. Structural comparisons of PIM1 to previously defined conformations of bacterial and human Lon reveal that the protease domain of PIM1 is in a proteolytically competent active state. In contrast to all prior structures, we observe an extended C-terminal tail forming an α-helix located along the base of the protease domain, engaged in hydrophobic interactions with the base of the core protease domain. Finally, despite structural conservation of the regions that flank the 50-residue charged insertion unique to *S. cerevisiae*, we do not observed ordered density corresponding to the charged residues, indicating this region is independently flexible. Biochemical assays targeting this charged region demonstrate minimal effects on Lon protease function, indicating that the yeast-specific charged insertion is dispensable for its enzymatic function.

## Results

### Substrate-bound structure of PIM1 demonstrates hexameric assembly and highlights conserved structural elements

Although atomic structures of bacterial and human Lon define a hexameric assembly for Lon protease, initial negative stain electron microscopy and analytical centrifugation suggested that yeast Lon (PIM1) oligomerizes into a heptameric assembly^12^. Of note, PIM1 from *S. cerevisiae* is defined by charged insertion of approximately 50 residues between the ATPase and protease domains (**Supplementary Figure 1**). To define the oligomeric state of PIM1 and understand the structural role of this charged insertion, we aimed to determine a high-resolution structure of PIM1. We recombinantly expressed and purified mature wild-type *S. cerevisiae* PIM1 lacking the mitochondrial targeting sequence from bacteria, hereafter referred to as PIM1. After size-exclusion chromatography, PIM1 was incubated with saturating amounts of ATPγS (1 mM), a slowly hydrolyzing ATP analog, and plunge-frozen for investigation by cryo-electron microscopy (cryo-EM). Remarkably, data collection and image analysis yielded the structure of PIM1 in a hexameric, closed, substrate-translocating state at a reported resolution of ~3.2 Å. Notably, and in stark contrast to all prior cryo-EM studies of Lon proteases in other species^13,14,16–18^, we did not observe the presence of a fully ADP-bound substrate-free oligomer, typically defined by the AAA+ domains adopting an open, left-handed spiral organization. We note a high degree of background in our micrographs, which we believe to be monomeric PIM1 derived from biochemical instability of the hexamer or particle disruption at the air-water interface (**Supplementary Figure 2A**). Indeed, previous structural analyses of oligomeric PIM1 required cross-linking by glutaraldehyde and demonstrated nucleotide-dependent stability^12^. This leads us to speculate that, unlike other Lon proteases, yeast PIM1 only hexamerizes in the presence of substrate.

Consistent with the structures of bacterial and human Lon as well as other AAA+ proteases^13–15^, PIM1 reveals a hexameric assembly defined by a right-handed spiral configuration (**Figure 1A**). Residues 529 to 1133, comprising the N-terminal 3-helix bundle, ATPase, and protease domains, were well-resolved to enable model building and refinement. This hexameric PIM1 assembly is highly similar to that of bacterial and human Lon with ~0.9 Å root mean squared deviation (RMSD) in the ATPase domain and ~0.6 Å RMSD in the protease domain. Despite the inclusion of residues 181 to 528, which compose the N-terminal domain in our construct, we only observed blurred density corresponding to this region in 2D class averages (**Supplementary Figure 2B**), suggesting that the N terminal domain of PIM1 is flexible. Although PIM1 was not purified in the presence of exogenous substrate, we observe substrate density intercalated with the substrate-translocating pore loop residues of Lon, which we modeled as a nine residue poly-alanine chain (**Figure 1B**). Across AAA+ proteins, a conserved aromatic-hydrophobic motif found in pore loop residues directly mediates AAA+-dependent substrate unfolding and translocation^15^. In agreement, ATP-bound PIM1 subunits interact with substrate via the conserved pore loop 1 aromatic residue Y674 and hydrophobic residue I675 (**Figure 1B**). These conserved pore loop 1 residues engage unfolded substrate with a “pincer-like” grasp and, according to the hand-over-hand model of translocation, the energy conferred by ATP hydrolysis is used to shuttle unfolded substrate towards the proteolytic chamber for degradation.

**Figure 1.**
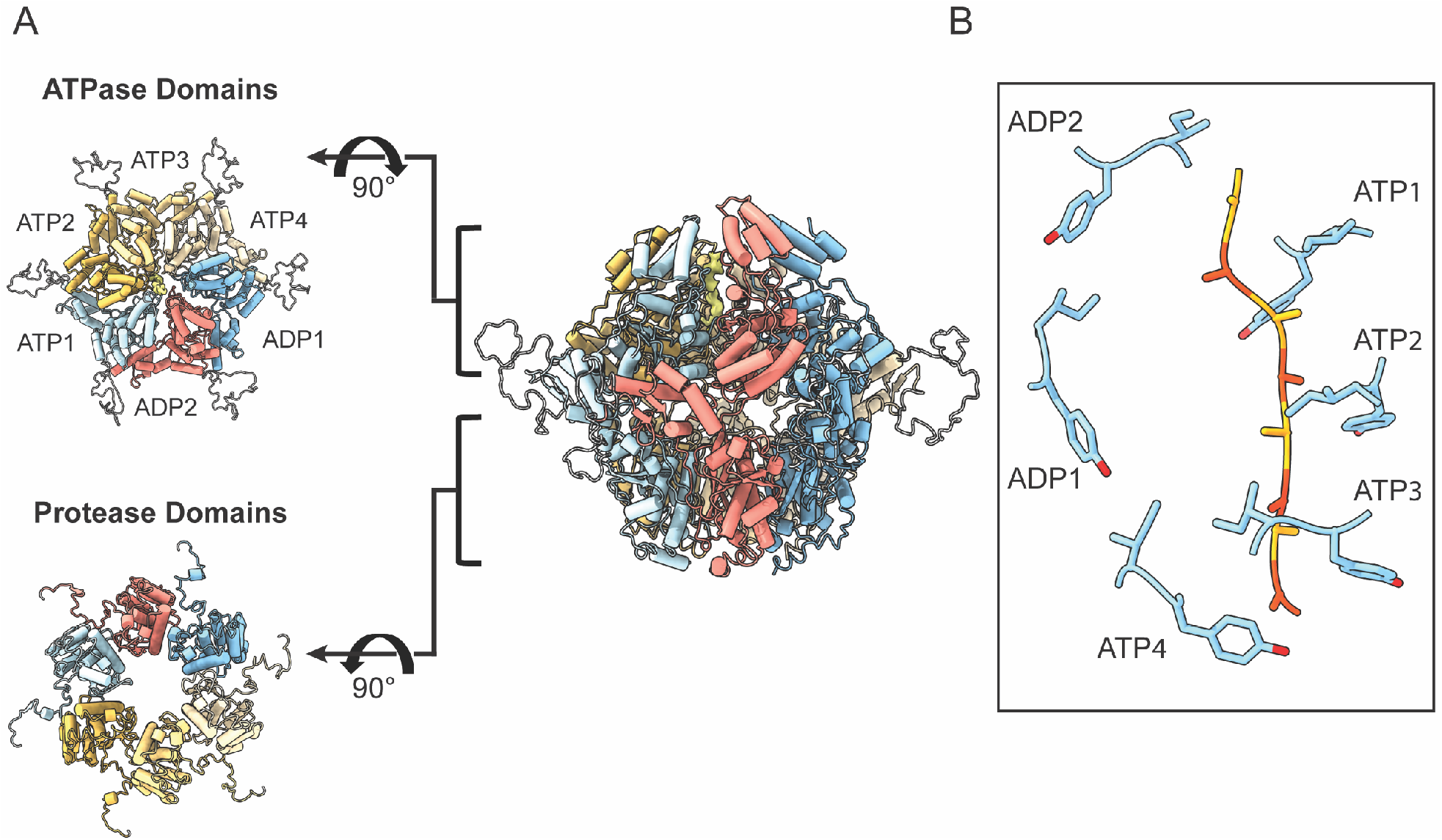
Architecture of substrate-translocating PIM1. **(A)** The substrate-translocating PIM1 atomic model (center) with orthogonal views of the ATPase (upper left) and protease (lower left) domains. PIM1 forms a closed, right-handed spiral staircase. Each subunit is designated a specific color based on its position in the right-handed spiral staircase and trapped substrate is colored in orange. **(B)** Conserved pore loop aromatic residue Y674 and hydrophobic residue I675 from the ATP-containing subunits engage unfolded substrate in a “pincer-like” grasp. Y675 and I675 are shown in stick representation. Density that corresponds to substrate is modeled as a polyalanine chain and colored in yellow and orange. Substrate-translocating PIM1 is characterized by four descending ATPase domains labeled ATP1-4 and two ascending “seam” subunits labeled ADP1 and ADP2.

Prior structural studies of substrate-translocating AAA+ proteins and Lon proteases are consistent with this model of hand-over-hand substrate translocation^13,14,16–18^. In this model, pore loop residues of ATP-bound subunits form a spiral arrangement wherein the lower-most subunit’s pore loop residue typically exists in an intermediary state. In concordance with the AAA+ organization of bacterial Lon proteases^13,18,19^, our substrate-translocating PIM1 structure consists of four descending ATPase domains (labeled here as ATP1-3, ATP4) and two ascending ‘seam’ subunits (ADP1-2). The ADP-bound subunits are found between the lowest and highest subunits of the substrate-bound spiral arrangement and ADP-bound subunit pore loop residues are indeed disengaged from substrate (**Figure 1B**, ADP1/ADP2). It is believed that as ATP hydrolysis occurs in the lowermost subunit ATP-bound subunit, this subunit releases its grip on substrate and the remaining 3 ATP-bound subunits drop to a lower register within the spiral staircase, translocating substrate a distance of two residues. Concurrently, the ADP2 subunit exchanges ADP for ATP and engages substrate at the top of the staircase^15^.

The nucleotide binding pockets in substrate-bound Lon protease are found in a cleft between the large and small subdomains of the ATPase domain. This cleft is located at a protomer-protomer interface and enables nucleotide-dependent changes to allosterically influence the nucleotide-bound subunit as well as its neighboring subunit. Structurally, the nucleotide-bound subunit is characterized by universally conserved structural motifs, such as the Walker A motif (important for nucleotide binding, K638 in PIM1) and the Walker B motif (important for nucleotide hydrolysis, E700 in PIM1), and partially conserved structural motifs, such as the sensor-1 residue (N750 in PIM1) and cis-acting sensor-2 residue (R820 in PIM1). These sensor-1 and cis-acting sensor-2 residues position the ATP β-γ phosphate bond for hydrolysis. Finally, a transacting arginine finger (R672) from the neighboring subunit interacts with the γ-phosphate to critically convey nucleotide status. The resolution of our reconstructed density enabled modeling of these nucleotide interactions (**Supplementary Figure 3**). The four subunits that directly engage substrate and form a continuous right-handed spiral staircase contain density consistent with an ATP molecule. These subunits are named ATP1, ATP2, ATP3, and ATP4 in order from the uppermost subunit of the ATPase staircase to the lowest. The remaining two subunits do not directly engage subunit and do not contain density for a γ-phosphate and are named ADP1 and ADP2.

### Substrate-translocating PIM1 protease domains are in a proteolytically competent configuration and contain a unique C-terminal extension necessary for protease activity

Upon translocation of substrate to the proteolytic chamber, PIM1 uses a serine-lysine catalytic dyad formed by S1015 and K1058 to cleave substrates. Previous structural examination of the bacterial Lon protease domain in substrate-free and substrate-bound conformations showed that in a substrate-free configuration, the S1015-containing helix folds into a 3-10 helix that sterically blocks access of substrate to the serine-lysine catalytic dyad^13,14^. Upon substrate-binding-induced symmetrization of the protease domain, this 3-10 helix unfolds to allow for a proteolytically competent serine-lysine catalytic dyad. The organization we observe in our PIM1 structure is consistent with the proteolytically competent conformation, where 3-10 helix is unfolded to form a substrate-binding groove and the serine-lysine dyad formed by S1015 and K1058 can be accessed by unfolded substrate (**Figure 2 A,B**). Previous structural work on the human Lon protease showed that substrate binding within the ATPase channel was insufficient to trigger a rearrangement of the protease domains to a symmetric proteolytically competent conformer, suggestive of a multi-level regulatory mechanism^14^. The symmetric organization of our PIM1 structure, which is consistent with bacterial structures, indicates that the human regulatory mechanisms are not conserved across eukaryotes.

**Figure 2.**
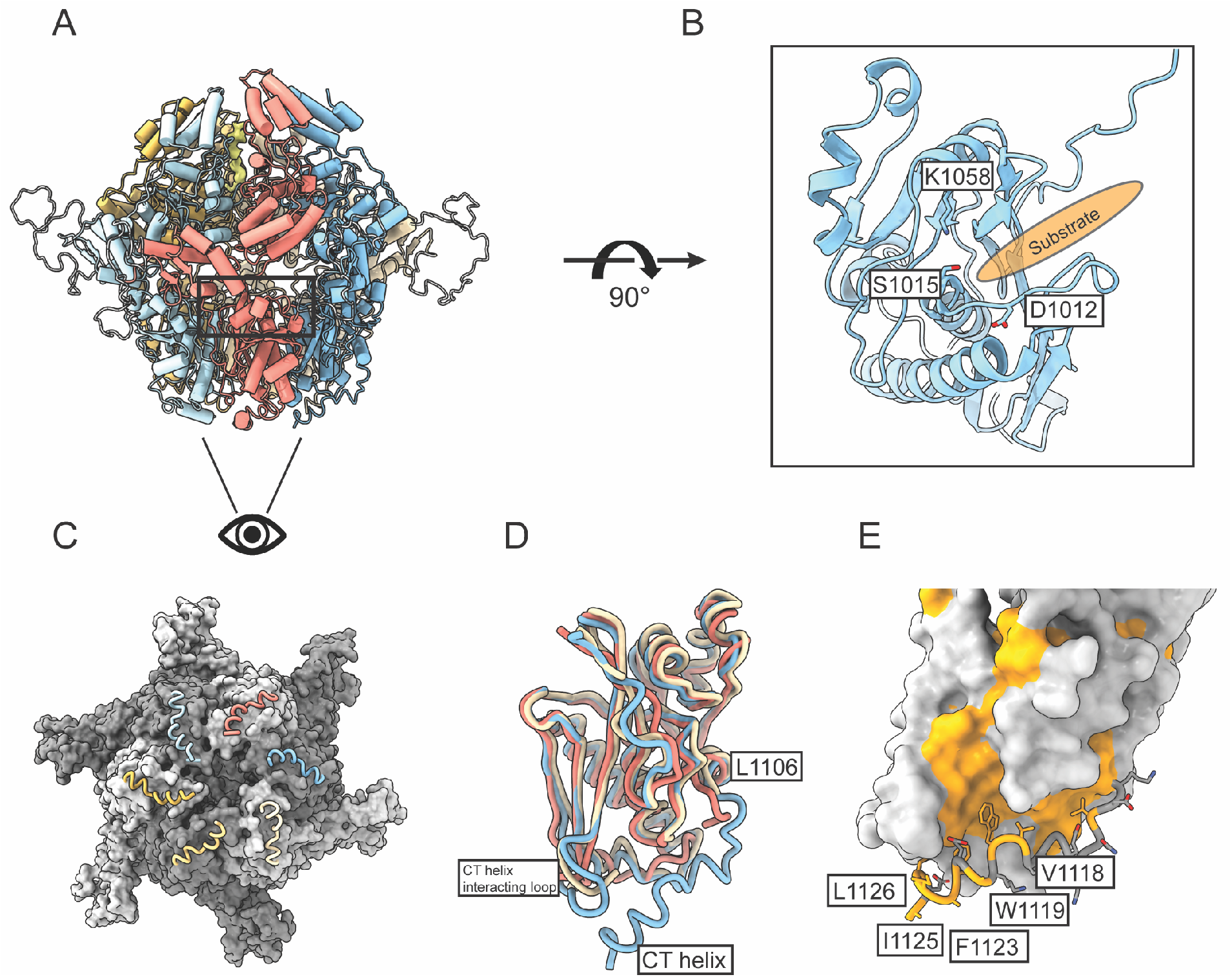
PIM1 protease domain is proteolytically competent and contains a unique C-terminal extension. **(A)** Atomic model of substrate-translocating PIM1. **(B)** Substrate-translocating PIM1 protease domains are proteolytically competent. The serine-lysine catalytic dyad is formed by S1015 and K1058. Upon substrate binding and symmetrization of the protease domain, a 3-10 helix that contains D1012 unfolds to enable formation of the proteolytically competent serine-lysine catalytic dyad. The substrate-binding groove of PIM1 depicted as orange oval. **(C)** Orthogonal view of the protease domain highlighting the PIM1-specific C-terminal extension. PIM1 is represented as a gray surface and the C-terminal extension is highlighted in worm representation. These C-terminal residues lie along the base of the protease domain and form intra-protomeric contacts. **(D)** Superposition of the PIM1 protease domain with that of substrate-translocating Yersenia pestis Lon (PDB ID: 6on2) and human Lon (PDB ID: 7krz) highlights Lon protease C-terminal residues and a PIM1-specific alpha-helix. Reconstructions of Lon protease C-termini to date have enabled confident model building to approximately the alpha-helix that ends at L1106 in PIM1. K1120 in PIM1 is the C-terminal-most conserved residue across Lon proteases according to sequence alignment (Supplementary Figure 1). **(E)** Examination of the C-terminal residues reveals that the C-termini are held in place by hydrophobic interactions between V1118, W1119, F1123, I1125, and L1126 of the C-terminus, F994 and A1028 of the core protease domain, and L952 and H953 of a loop at the base of the core protease domain. PIM1 is depicted as a surface representation with hydrophobic residues colored in orange. The C-terminal helix and associated hydrophobic residues are highlighted in worm and stick representation, respectively.

PIM1 contains a C-terminal extension that is not present in bacterial or human Lon. This C-terminal extension was unexpectedly well-resolved as an α-helix in our reconstruction, and runs along the base of the protease domain (**Figure 2C**). Structural comparison of the PIM1 protease domain with that of substrate-translocating *Yersenia pestis* Lon (PDB ID: 6on2) and human Lon (PDB ID: 7krz) highlights structural conservation up to L1106 of PIM1 (**Figure 2D**). Although sequence alignment of PIM1 with *Yersinia pestis* Lon and human Lon suggest that these protease domains are homologous up to K1120 of PIM1 (**Supplementary Fig 1**), residues beyond the structurally conserved α-helix in bacterial or human Lon could not be modeled due to disorder of the C-terminal regions. In PIM1, the conserved α-helix is followed by an ordered loop extending to K1120, and ending with an α-helix (A1121 to K1129) that is unique to PIM1. Situated at the base of the protease domain, the C-terminal helix is stabilized by intra-subunit hydrophobic interactions among residues V1118, W1119, F1123, I1125, and L1126 of the C-terminus, residues F994 and A1028 of the core protease domain, and residues L952 and H953 of a loop at the base of the core protease domain, hereafter referred to as the CT helix-interacting loop (**Figure 2E**).

To define the functional importance of the yeast-specific C-terminal extension, we first inspected available structures of substrate-free and substrate-translocating Lon proteases. Structural analyses revealed that regions around residues F994 and A1028 of the core protease domain and the CT helix-interacting loop are structurally invariant in both substrate-free and substrate-translocating conformations. Based on available structural information, the functional importance of the yeast-specific C-terminal extension is obscure. Given the location of the C-terminal helix at the base of the protease chamber, we speculate that the C-terminal helix may be involved in cleaved peptide release, although further investigation of this potential activity is necessary.

### PIM1 Charged Insertion Appears Structurally and Biochemically Dispensable for Proteolytic Function

One of the aims of our study was to understand the structural importance of the charged insertion of approximately 50 residues between the ATPase and protease domains of PIM1. Unexpectedly, in our reconstruction of substrate-translocating PIM1, we did not observe density for this yeast-specific charged insertion. We observe strong density preceding and subsequent to the charged insertion at K840 and N898, respectively. Both the K840-containing α-helix and the N898-containing β-strand are structurally conserved elements across the ATPase domain of Lon orthologues (**Figure 3A**). Based on the apparent disorder of the charged insertion in our structure of PIM1, we wondered whether the insertion is functionally dispensable for PIM1 enzymatic activities.

**Figure 3.**
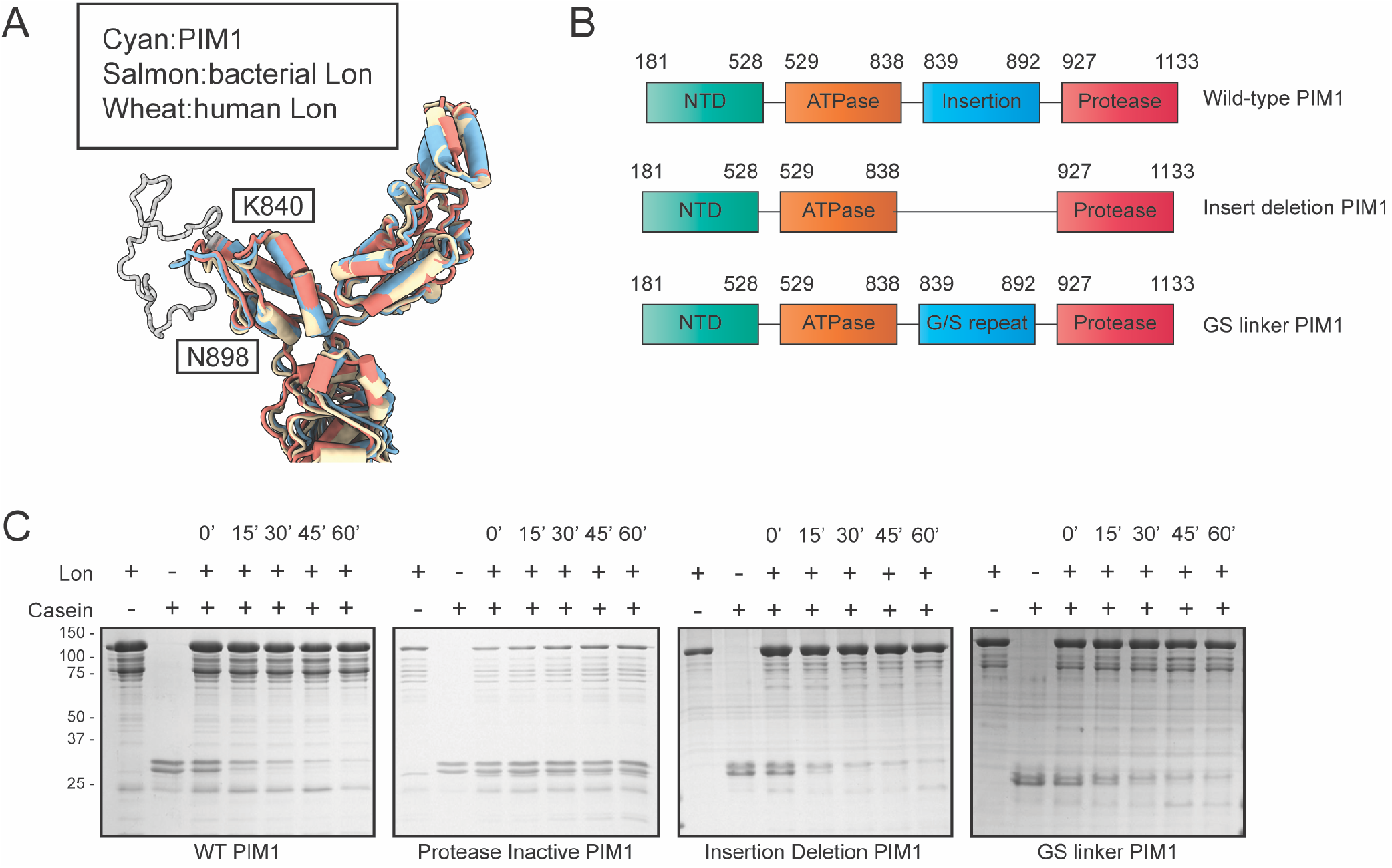
PIM1-specific charged insertion appears structurally and biochemically dispensable for proteolytic function. **(A)** Superposition of PIM1 ATPase domains with that of substrate-translocating Yersenia pestis Lon (PDB ID: 6on2) and human Lon (PDB ID: 7krz) highlights strong structural conservation. The K840-containing alpha-helix and the N898-containg beta-strand that precede and conclude the yeast-specific charged insertion are structurally conserved elements across ATPase domains of Lon orthologues. **(B)** Cartoon diagram of mutant PIM1 constructs used to test the biochemical importance of the charged insertion. PIM1 structural domains and associated residues are specified. **(C)** Gel-based proteolysis assay of mutant PIM1 constructs using casein as a model substrate over a one-hour time course. PIM1 is 109 kDa and casein is observed as a doublet band around 30 kDa.

To clarify the biochemical implications of this charged insertion on PIM1 function, we generated two mutants: 1) charged insertion deletion (PIM1-INSdel); 2) replacement of charged insertion with a glycine-serine linker of identical length (PIM1-GSlink) (**Figure 3B**). With the PIM1-INSdel construct, we aimed to examine whether the charged insertion was completely dispensable; with the PIM1-GSlink construct, we aimed to discriminate whether it was the amino acid composition or insertion length that was critical for PIM1 function.

For both PIM1-INSdel and PIM1-Gslink mutants, we observed close to wild-type levels of ATP-dependent substrate degradation in a gel-based proteolysis assay using casein as a model substrate and a protease-inactive PIM1 as a control (**Figure 3C**). This demonstrates that this charged insertion is dispensable for Lon proteolytic activity. Based on our biochemistry and structural anaylses, we speculate that this yeast-specific insertion may be required for recruitment of substrates or PIM1 adaptor proteins.

## Discussion

Here, we determined the cryo-EM structure of substrate-translocating PIM1 from *Saccharomyces cerevisiae* to demonstrate that PIM1 forms a hexameric assembly, consistent with multiple Lon protease structures from bacteria and human. As observed in other AAA+ protein translocases, aromatic pore loop residues directly bind substrate at the central channel of the ATPase domain to processively shuttle unfolded polypeptides in a hand-over-hand mechanism coordinated with ATP binding and hydrolysis.

Examination of the PIM1 protease domains reveals that they organize into a C6-symmetric assembly that is proteolytically competent, similar to the bacterial form and the human Lon when bound to bortezomib^13,14^. The reconstruction also shows the presence of an extended, ordered C-terminus, which includes an α-helix located along the base of the protease domain. Structurally, the role of these yeast-specific C-terminal residues is ambiguous, as the central cavity is not noticeably occluded by this 14-residue extension and interactions of the C-termini appear to be strictly intra-protomeric. We speculate that these yeast-specific C-terminal residues could facilitate cleaved peptide release, which warrants future investigation.

Of note, PIM1 from *S. cerevisiae* is defined by a charged insertion of approximately 50 residues between the ATPase and protease domains. Although our reconstruction confirms structural conservation of the regions before and after this charged insertion, we do not observe density for the charged insertion loop itself. To understand the functional role of the charged insertion, we examined Lon mutants that delete the insertion and replace the insertion with a non-specific glycine-serine linker of identical length. In biochemical assays examining protease activity, we demonstrate minimal effects on Lon protease function. Thus, we propose that the yeast-specific charged insertion is dispensable for enzymatic function.

Our investigation of PIM1 suggests that Lon proteases are structurally conserved across all kingdoms of life. Indeed, previous work replacing PIM1 with a mitochondrially targeted Lon protease from *Escherichia coli* has demonstrated that Lon proteases are highly functionally redundant^20^. In this previous study, Teichmann and colleagues demonstrated that wild-type *E.coli* Lon, but not protease-inactive Lon could rescue respiration deficiency and mitochondrial DNA integrity of PIM1 knock-out cells at 30 degrees C and mediate degradation of mitochondrial matrix proteins. Notably, at 37 °C, wild-type *E.coli* Lon could not rescue respiration deficiency and mitochondrial DNA integrity of PIM1 knock-out cells. Teichmann and colleagues speculated that this could be a consequence of non-overlapping activities in protein domains that are not conserved between mitochondrial and prokaryotic homologues.

Previous work using Lon protease from α-proteobacteria *Caulobacter crescentus* has suggested that a positively charged patch on the surface of Lon protease ATPase domains directly mediates binding to DNA^21^. Here, mutation of four lysines to glutamic acid abrogates the capacity of *C. crescentus* Lon to bind DNA. These four lysines lie on the lateral surface of the ATPase domain and it is tempting to speculate that the charged residues that make up the yeast-specific insertion would sterically facilitate or interfere with binding. Recent work has also demonstrated that PIM1 modulates mitochondrial DNA copy number through direct interactions with Mrx6, an understudied protein characterized by a PET20 domain^22^. Here, we speculate that an unstructured region, such as that of the PIM1 charged insertion, may serve as handle for binding by yeast-specific binding partners or DNA, for which further investigation is necessary. For disordered domains and proteins at large, ligand binding can confer dramatic changes in structural dynamics. Our structure of PIM1 in a substrate-translocating state provides a starting point to define the functional implications of the yeast-specific charged insertion.

## Materials and Methods

### Protein expression and purification

The wild-type (WT) PIM1 construct without the mitochondrial signal sequence (Met182 – Asp1133) was cloned into a modified pET41a expression vector containing a 6x-His tag at the N-terminus. The Q5 site-directed mutagenesis kit from New England Biolabs was used to generate the protease inactive S1015A mutant. E. coli strain BL21 (DE3) Codon Plus RIPL (Agilent Technologies) was used for recombinant protein expression. The transformed BL21 RIPL cells were grown at 37 °C until the OD reached 0.6~0.8. Protein expression was induced at 16 °C overnight with 0.5 mM IPTG. Cells were harvested by centrifuge at 30,000 g for 20 minutes at 4 °C. The cell pellets were resuspended and lysed using sonication in a lysis buffer containing 25 mM Tris-HCl (pH 8.0), 500mM NaCl, 10% glycerol. The lysis was cleared by ultra-centrifuge followed by Ni-NTA affinity binding. The protein was washed and eluted by wash buffer (lysis buffer supplemented with 50 mM imidazole), and elution buffer (lysis buffer supplemented with 300 mM imidazole). PIM1 was further purified through a Superose 6 size-exclusion chromatography (SEC) column in a buffer containing 25mM Tris-HCl (pH 8.0), 150mM NaCl, 5mM MgCl_2_, 5 mM DTT. Peak fractions containing hexameric PIM1 were pooled, concentrated and flash-frozen for storage at −80 °C. Protease inactive PIM1 and PIM1 mutants were expressed and purified similarly.

### Sample preparation for electron microscopy

3 mg/ml WT PIM1 was incubated with 1mM ATPγS (MilliporeSigma) on ice for 20 minutes. 4μl of the sample was applied onto 300 mesh R1.2/1.3 UltraAuFoil Holey Gold grids (Quantifoil) that were plasma cleaned for 15 s using Pelco glow discharge cleaning system (TED PELLA, INC.) with atmospheric gases at 15 mA. The sample was plunge-frozen using a Thermo Fisher Vitrobot Mark IV at 4 °C, 5 s, 100% humidity, and a blot force of 0 using Whatman #1 blotting paper.

### *In vitro* ATP hydrolysis assay

*In vitro* ATP hydrolysis was carried out in LONP1 activity buffer (50 mM Tris-HCl pH 8, 100 mM KCl, 10 mM MgCl_2_, 10% Glycerol, and 1 mM DTT) in the presence or absence of 2.5 mM ATP. Casein (9.7 μM) was added to the reaction mixture when measuring substrate-induced ATPase activity. Reaction components and purified LONP1 (0.1 μM) were incubated separately at 37°C for five minutes. LONP1 was added to initiate ATP hydrolysis and the amount of free inorganic phosphate at each timepoint was measured by adding Malachite Green Working Reagent (Sigma-Aldrich) and incubating components for 30 minutes at room temperature for color development before measuring absorbance at 620 nm (OD_620_). Three biological replicates were performed for each LONP1 mutant in the presence and absence of substrate. Data were fit to a straight line and the slope were extracted to calculate ATP hydrolysis rates. GraphPad Prism software was used for data analysis. Mean and standard error of mean (SEM) was calculated by performing column statistics. ANOVA was used to calculate p-value, as indicated.

### *In vitro* proteolysis assay

Each reaction was carried out in an activity buffer composed of 50 mM Tris-HCl (pH 8.0), 100 mM KCl, 10 mM MgCl_2_, 10% glycerol, and 5 mM ATP. 0.1 mg/ml of purified PIM1 and PIM1 mutants were incubated at 37 °C with or without 0.08 mg/ml beta-casein (Sigma-Aldrich). Aliquots were taken at specified time intervals and immediately mixed with 6X SDS sample buffer to terminate the reaction. Reaction products were resolved by gel electrophoresis.

### Cryo-EM data collection and processing

Cryo-EM data were collected on a Thermo-Fisher Talos Arctica transmission electron microscope operating at 200 keV using parallel illumination conditions^23^. Micrographs were acquired with a Gatan K2 Summit direct electron detector with a total electron exposure of 50e^−^/Å ^2^ as a 97-frame dose-fractionated movie during a 9.7 s exposure time. The Leginon data collection software^24^ was used to collect 1,394 micrographs at 36,000 nominal magnification (1.15 Å /pixel) with a nominal defocus ranging from −0.8 μm to −1.5 μm. Stage movement was used to target the center of four 1.2 μm holes for focusing, and image shift was used to acquire high magnification images. The Appion image processing wrapper^25^ was used to run MotionCor2 for micrograph frame alignment and dose-weighting in real-time during data collection^26^. All subsequent image processing was done in cryoSPARC^27^. The CTF parameters were estimated using Patch CTF estimation (multi). Representative views of 2D classes from negative-staining data were used as templates for template-based particles picking in cryoSPARC, which resulting in picked 497,192 picked particles. Particle coordinates were extracted at 1.15 Å /pixel from the motion-corrected and dose-weighted micrographs with a box size of 256 pixels. 100 classes of 2D were classified using default parameters. 2D classes (111,203 particles) displaying high-resolution secondary features were selected and the 111,203 particles belonging to those classes were used to generate a reference-free 3D Ab-initio model. The selected particles were further classified with 3D classification into 3 classes using default parameters. Class 1 (41,268 particles) and class 2 (40,675 particles), which displayed high-resolution structures details, were selected for CTF refinement followed by 3D NU-Refinement, which resulted in the final reconstruction with reported resolution of ~3.21 Å according to a Fourier Shell Correlation (FSC) cutoff at 0.143 (**Supplementary Figure 2**).

### Atomic model building and refinement

A homology model of substrate-translocating PIM1 was generated with SWISS-MODEL, using a previously solved cryo-EM structure of human LONP1 (PDB ID: 7KSM) as a starting model^14,28^. The resulting model was initially docked into the PIM1 reconstructed density map using UCSF Chimera. Real-space structural refinement and manual model building were performed with Phenix^29^ and Coot^30^, respectively. Phenix real-space refinement includes global minimization, rigid body, local grid search, atomic displacement parameters, and morphing for the first cycle. It was run for 100 iterations, 5 macro cycles, with a target bonds RMSD of 0.01 and a target angles RMSD of 1.0. The refinement settings also include the secondary structure restraints, Ramachandran restraints. Figures for publication were prepared using PyMol, UCSF Chimera, and UCSF ChimeraX^31,32^.

## Data availability

The reconstructed density map and atomic model of PIM1 have been deposited at the Electron Microscopy Data Bank and PDB under accession numbers EMDB: EMD-25505 and PDB: 7XSO, respectively. All data needed to evaluate the conclusions in the paper are present in the paper and the supporting information.

## Acknowledgements

We thank J.C. Ducom at Scripps Research High Performance Computing and C. Bowman at Scripps Research for computational support, and B. Anderson at the Scripps Research Electron Microscopy Facility for microscopy support. This work is supported by the NIH (NS095892 to R.L.W. and G.C.L). A.S.S is supported by the NIH F31AG071162 and the Olson-King Endowed Skaggs Fellowship from The Scripps Research Institute. Computational analyses of EM data were performed using shared instrumentation funded by NIH S10OD021634 to G.C.L.

## Supplementary Materials

**Supplementary Figure 1:**
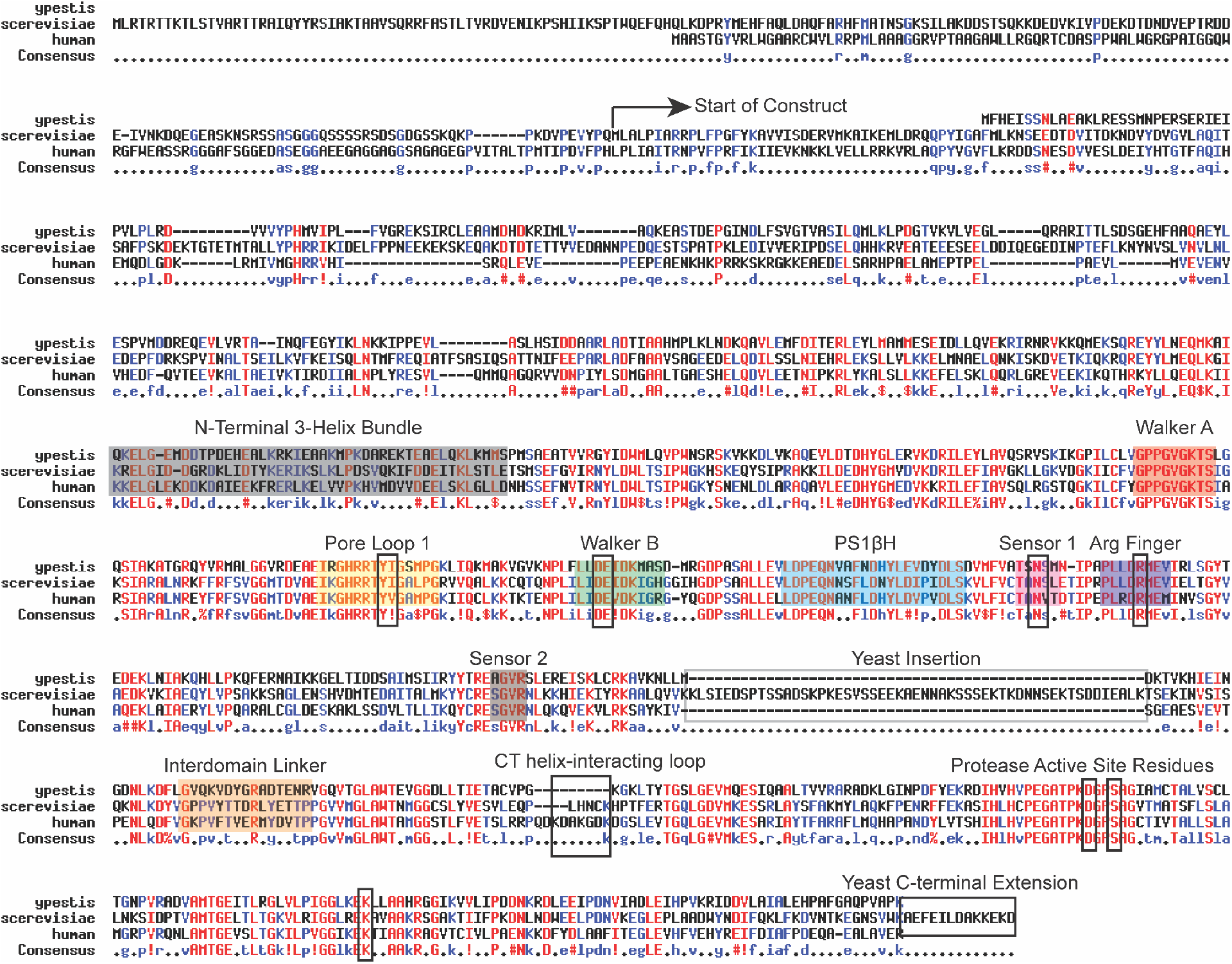
Sequence alignment of *S. cerevisiae* Lon (PIM1), *Y. pestis* Lon, and human Lon. Sequence alignment highlights conserved structural elements: N-terminal 3-helix bundle (gray), Walker A (orange), Pore loop 1 (yellow, aromatic residue Y674 and hydrophobic residue I675 boxed), Walker B (green, nucleotide coordinating acidic residues boxed), Pre-sensor 1 beta-hairpin (light blue), Sensor 1 (pink, N750 boxed), Arginine finger (purple, R762 boxed), Sensor 2 (gray), yeast-specific insertion (gray box), interdomain linker (yellow), CT helix interacting loop (black box), protease active sites (black box, S1015; K1058; D1012), Yeast-specific C-terminal extension.

**Supplementary Figure 2:**
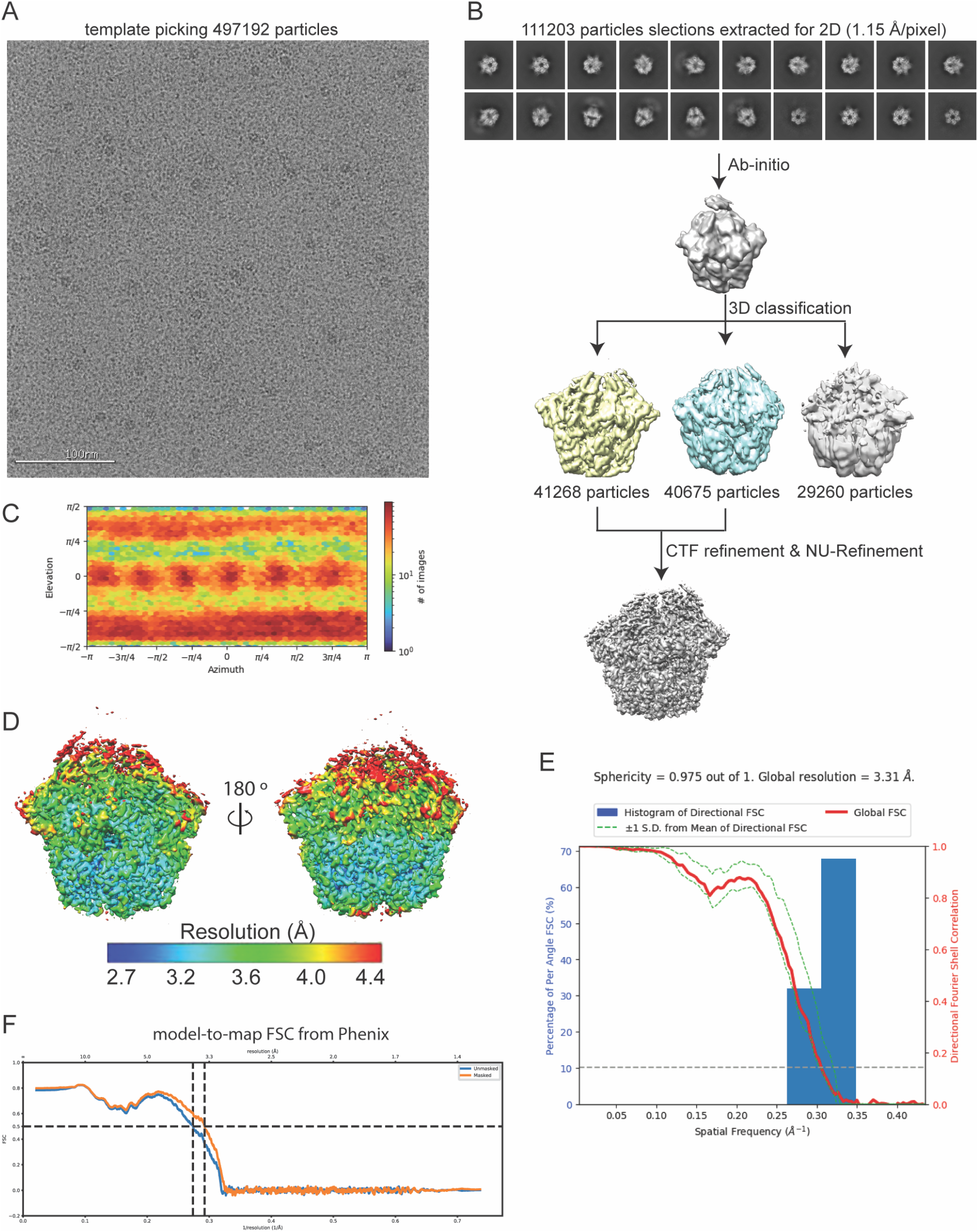
Cryo-EM structure determination of PIM1. **(A)** Representative micrograph of cryo-EM data collection, showing high background of what appears to be monomeric PIM1. **(B)** Workflow of cryo-EM data processing using cryoSPARC software^27^. The final 3D reconstruction map was used for model building and refinement. **(C)** Euler angle distribution plot of the particles used in the final reconstruction. **(D)** Final reconstruction filtered and colored by local resolution from cryoSPARC. **(E)** 3-Dimensional Fourier Shell Correlation (3DFSC)^33^ of the final reconstruction reporting a global resolution of 3.3 Å. **(F)** Model-to-map FSC plot from Phenix validation.

**Supplementary Figure 3:**
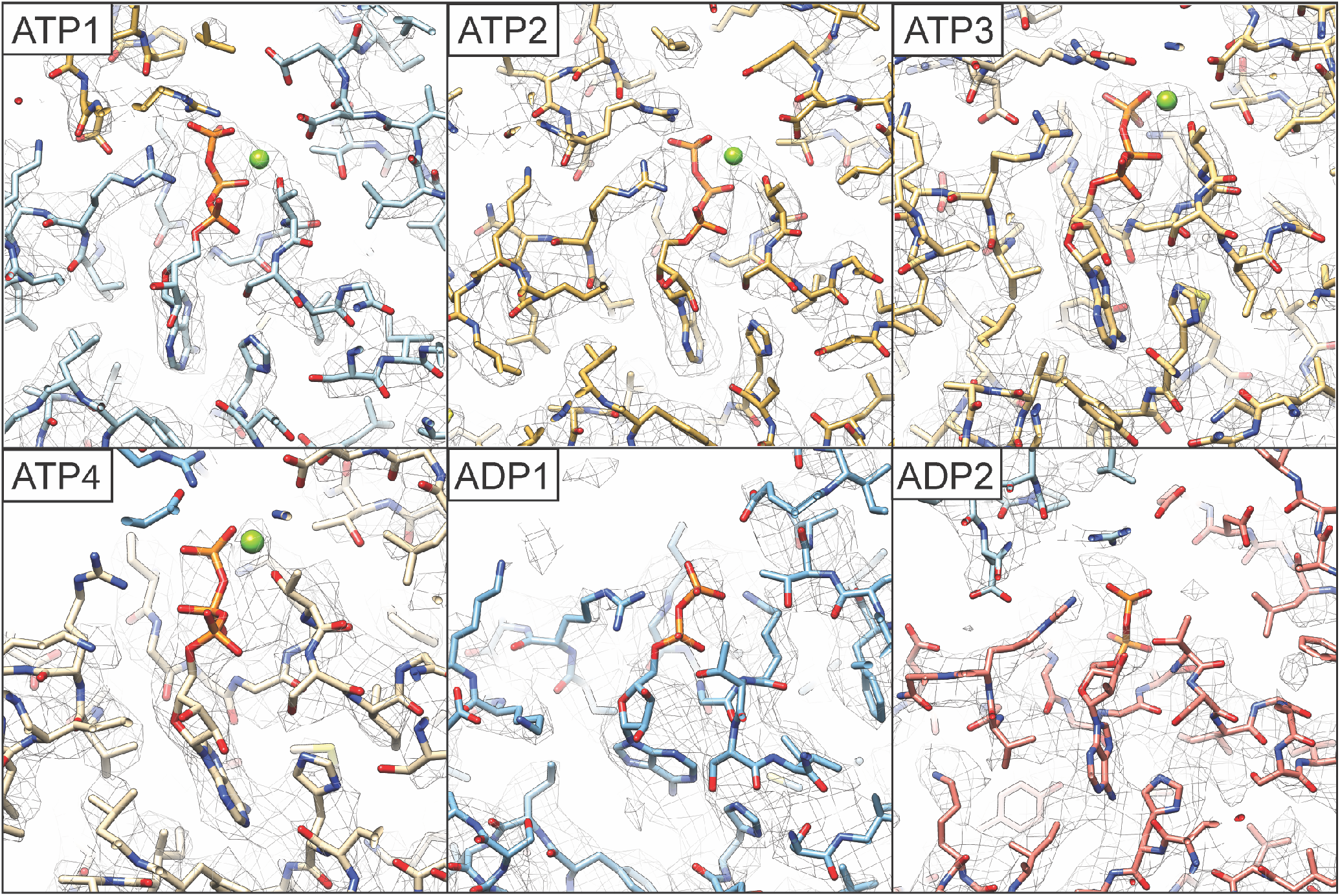
Reconstructed density for nucleotide binding pockets enables assignment of nucleotide states. Views of the nucleotide binding pocket of all six PIM1 subunits colored according to Figure 1, with atomic model represented as sticks and EM density as a mesh.

**Table S1.** Cryo-EM data collection, refinement, and validation statistics

|  |  |
| --- | --- |
| EMDB ID | 25502 |
| PDB ID | 7SXO |
| **Data Collection** |  |
| Microscope | Talos Arctica |
| Camera | K2 Summit |
| Magnification (nominal/at detector) | 36,000/43,478 |
| Voltage (kV) | 200 |
| Data acquisition software | Leginon<sup>24</sup> |
| Exposure navigation | Image shift to 16 holes |
| Total electron exposure (e<sup>-</sup>/Å<sup>2</sup>) | 50 |
| Exposure rate (e<sup>-</sup>/pixel/sec) | 6.8 |
| Frame length (ms) | 100 |
| Number of frames | 97 |
| Pixel size (Å/pixel)* | 1.15 |
| Defocus range (μm) | 0.7 to 1.5 |
| Micrographs collected (no.) | 1,394 |
| **Reconstruction** |  |
| Total extracted picks (no.) | 497,129 |
| Particles used for 3D (no.) | 111,203 |
| Final Particles (no.) | 111,203 |
| Symmetry | C1 |
| Resolution (global) (Å) |  |
| FSC 0.5 (unmasked / masked) | 4.4 / 3.5 |
| FSC 0.143 (unmasked / masked) | 3.8 / 3.2 |
| Local resolution range (Å) | 3.2 - 5.0 |
| 3DFSC Sphericity<sup>62</sup> (%) | 97.5 |
| Map sharpening B-factor (Å<sup>2</sup>) | -82.7 |
| **Model composition** |  |
| Protein residues | 3,613 |
| Ligands | 12 |
| **Model Refinement\*** |  |
| Refinement package | Phenix<sup>29</sup> |
| MapCC (volume/mask) | 0.75/0.74 |
| R.m.s. deviations |  |
| Bond lengths (Å) | 0.003 |
| Bond angles (°) | 0.619 |
| **Validation** |  |
| Map-to-model FSC 0.5 | 3.4 |
| Ramachandran plot |  |
| Favored (%) | 93.04 |
| Allowed (%) | 6.96 |
| Outliers (%) | 0.0 |
| MolProbity score | 1.94 |
| Clashscore | 9.2 |
| Poor rotamers (%) | 0.0 |
| C-beta deviations | 0.0 |
| CaBLAM outliers (%) | 4.85 |
| EMRinger score<sup>34</sup> | 2.05 |

